# *ankrd1a* consistently marks cardiomyocytes bordering the injury or scar area and affects their dedifferentiation during zebrafish heart regeneration after cryoinjury

**DOI:** 10.1101/2025.06.24.661322

**Authors:** Srdjan Boskovic, Mirjana Novkovic, Andjela Milicevic, Emilija Milosevic, Jovana Jasnic, Mina Milovanovic, Snezana Kojic

## Abstract

In contrast to humans, zebrafish have a remarkable ability to regenerate injured heart through a highly orchestrated process involving all cardiac structures. To replace the lost myocardium, resident cardiomyocytes (CMs) dedifferentiate and proliferate, invading the injured area. The response of the myocardium is preceded by the activation of the epicardium and endocardium, which form active scaffolds to provide mechanical and paracrine support to guide regeneration. New CMs use protrusions to migrate and invade fibrotic injured tissue, to replace it with functional myocardium. Here, we investigated the expression profile of the stress-responsive *ankrd1a* gene in different cardiac structures, at key time points during regeneration, aiming to gain insight into its precise roles during zebrafish heart regeneration. In the *TgBAC(ankrd1a:EGFP)* reporter line, transgene upregulation was restricted to the myocardium, initiated as early as 15 hours post-cryoinjury, and consistently marked CMs bordering the injury or scar area during regeneration. Transcriptome profiling and immunostaining revealed a potential role of *ankrd1a* in regulating CMs’ dedifferentiation, as well as changes in expression of genes associated with antigen presentation and extracellular matrix composition in the *ankrd1a* mutant. Our results indicate that the *ankrd1a* is dispensable for ventricle regeneration after cryoinjury and may be considered as a marker and fine-tuner in the healing process of injured cardiac muscle.

## INTRODUCTION

Among the many advantages that the zebrafish offers as a model system is the opportunity to study the regeneration of many organs, including the heart (1). Zebrafish regenerate their hearts after several kinds of injury, such as cryoinjury, ventricle resection, and ablation of cardiomyocytes (CMs) (1–5). Among these models, ventricular cryoinjury, followed by inflammation and massive apoptosis of cells affected by the injury, most closely resembles the processes in mammalian myocardial infarction and demonstrates the robust regenerative capacity of zebrafish (3). During cardiac regeneration, several partially overlapping processes take place: inflammatory response to cell necrosis and removal of dead cells, formation of a fibrin clot that closes the wound and production of collagen by fibroblasts, and formation of new myocardium (1–6). New CMs are derived from pre-existing CMs (7) that after dedifferentiation, proliferation, and redifferentiation (7, 8), use protrusions to migrate and invade injured fibrotic tissue to gradually replace it with functional myocardium (9). In most cases, complete regeneration occurs within 60 days (3), while damaged heart muscle regains its function within 90 days (10).

Zebrafish heart regeneration is a complex and precisely orchestrated process which requires the integration of a multicellular response (11). It involves all layers of the heart and numerous cell types, including CMs, epicardial and endocardial cells, immune cells, and connective tissue cells (12). The most intriguing processes in zebrafish heart regeneration upon cryoinjury occur in the region between injured and uninjured tissue, the so-called border zone (13). In this zone, the number of proliferating CMs, which contribute to myocardium regeneration, is the highest and their proliferation peak is at 7 days after cryoinjury (dpci). To become prone to proliferation, CMs of the border zone and under the epicardium dedifferentiate. This process occurs by the sarcomere rearrangements, reduced expression of mature sarcomere proteins, increased expression of developmental myosins, changes in intercellular connections, and re-entry into the cell cycle (7, 8).

Epicardium and endocardium support the proliferation of existing CMs at the injury border. Epicardium is required for zebrafish heart regeneration by exerting paracrine function and inducing the re-expression of developmental signalling markers (14). Following cardiac injury, activated epicardial cells induce stimulators of CMs’ cell cycle re-entry and proliferation (4, 15–18). Cells of epicardial origin migrate to and accumulate in the wound, forming a thick epicardial cap over the injury site and depositing extracellular matrix (ECM) components such as fibronectin and collagens to modulate the extracellular environment for optimal regeneration (19–21). Similarly, endocardium contributes to the structure and regulation of the ECM in the regenerating heart (22, 23). As a mechanosensitive tissue, endocardium sends signals to the myocardium in response to disturbed blood flow caused by cardiac injury. After the induction of epicardial response, the injury provokes the endocardium to form a cohesive regenerating sheet on the inside of the wound (22). Endocardial cells proliferate (with a peak at 3 dpci) and undergo morphological changes directed towards the acquisition of a motile phenotype (18, 22). They also upregulate the expression of collagens and deposit them in scar tissue (22, 23).

Cardiac regeneration occurs following substantial stress impacting the heart. A significant role in regulating cellular responses to stress, particularly in cardiac tissue, is played by cardiac ankyrin repeat domain protein ANKRD1 (or CARP), which belongs to the conserved family of stress-inducible proteins, the muscle ankyrin repeat proteins family (MARPs) (24–26). ANKRD1 is detected within the sarcomeric I-band, which serves as a mechanical controlling hub of CMs’ ability to relax. It is a notable regulator of the titin-associated stretch-sensing machinery via interaction with sarcomeric proteins titin (27, 28) and myopalladin (29), actin filament (27, 28) and intermediate filament component desmin (30). In the nucleus, ANKRD1 acts as a transcriptional cofactor, negatively regulating myocardial genes expression (31–33) In the human and rodent heart ANKRD1 is involved in development and molecular mechanisms of stress-induced remodelling (25, 31–35). Altered expression of *ANKRD1* has been linked with various pathological conditions affecting the heart (25, 36). When overexpressed, it contributes to heart failure (37), and progressive diastolic dysfunction in mice (38). On the other hand, genetic ablation of *ANKRD1* prevents the development of dilated cardiomyopathy phenotype (39) and attenuates chemically induced cardiac hypertrophy (40). Moreover, several *ANKRD1* mutations have been identified in patients with hypertrophic and dilated cardiomyopathy and diastolic dysfunction (24, 36). In zebrafish heart, *ANKRD1* orthologue, *ankrd1a* was found significantly upregulated upon injury and increased physical activity, suggesting a conserved stress response function (41, 42).

We have previously reported upregulation of *ankrd1a* in regenerating zebrafish heart, more specifically in CMs of the border zone (42). In this study we expanded our investigation of the *ankrd1a* expression profile using *TgBAC*(*ankrd1a:EGFP*) reporter line and show that transgene upregulation is restricted to the myocardium and initiates as early as 15 hours post-cryoinjury (hpci). Transgene expression consistently marks CMs bordering injury or scar area during regeneration. Transcriptome profiling and immunostaining revealed a potential role of *ankrd1a* in regulating CMs dedifferentiation, as well as changes in expression of genes associated with antigen presentation and extracellular matrix composition in *ankrd1a* mutant. Despite this, our results indicate that *ankrd1a* function is dispensable for ventricle regeneration after cryoinjury.

## MATERIAL AND METHODS

### Zebrafish husbandry and transgenic lines

Zebrafish were maintained according to standard protocols (https://zfin.org) under 14h light/10h dark cycle at 28°C. Animals were anesthetized with a buffered solution of 0.6 mM tricaine (MS-222 ethyl-m-aminobenzoate, Sigma-Aldrich). Cryoinjury of the heart and sham operation were performed as previously described (15). The experimental groups consisted of randomly assigned, age-matched animals of both genders, ranging in age from 4 to 12 months. The number of zebrafish in each experiment is given in figure legends. Husbandry and procedures were performed following institutional and national ethical and animal welfare guidelines, harmonised with Directive 2010/63/EU (amended by Directive 2024/1262). Experiments were approved by the Veterinary Directorate, Ministry of Agriculture, Forestry and Water Economy of the Republic of Serbia (323-07-00771/2021-05 and 000094465/2024). Transgenic zebrafish lines used in this study include *TgBAC*(*ankrd1a*:*EGFP*)^bns505^ (42), *Tg*(-0.8*myl7*:*nls-DsRedExpress*)^hsc4^ (43), *Tg*(*kdrl*:*RAS-mCherry*)^s916^ (44) and *ankrd1a*^bns502^ (45).

### Tissue fixation and sectioning

For immunofluorescence and histological staining, hearts were extracted, fixed in 4% paraformaldehyde (PFA) in phosphate buffer saline (PBS) for 1h at room temperature (RT), and cryopreserved with 30% sucrose in PBS at 4°C for 12-16h. After embedding into a frozen section medium (Killik, Bio Optica), hearts were stored at −80°C. Ten- or 60 µm cryosections were prepared using CM 1860 UV cryostat (Leica). Tissue sections were collected on Superfrost Plus Microscope slides (Thermo Fisher Scientific), air-dried, and stored at -20°C.

### Immunofluorescence and histological staining

Immunofluorescence staining was performed as previously described for adult zebrafish heart sections (46). After initial washing with PBST (PBS, 0,1% Tween 20) and water, samples were permeabilized with 3% H_2_O_2_ in methanol for 1h at RT. Slides were then washed with water and PBST, and incubated in a blocking solution (2% horse serum, 0,2% Triton X-100, 1% DMSO in PBS) for 1-3h at RT. Incubation with primary antibodies diluted in blocking solution was performed overnight at 4°C. The following day, primary antibodies were removed by washing the slides with PBST for 30 minutes. Incubation with secondary antibodies diluted in blocking solution was done for 2h at RT. After a 30-minute final wash with PBST, the samples were mounted with fluorescence mounting medium (Dako, Agilent Technologies).

The following primary antibodies were used at the indicated dilutions: chicken anti-GFP (ab13970, Abcam, 1:750), mouse anti-embCMHC (N2.261, DSHB, 1:50), rabbit anti-DsRed (632496, Takara Bio, 1:1000), rabbit anti-Caveolin-1 (3267, Cell Signaling, 1:600), mouse anti-PCNA (ab29, Abcam, 1:250). The following Alexa Fluor^TM^ secondary antibodies were used: anti-chicken AF488 (A32931), anti-mouse AF647 (A-21236), anti-rabbit AF647 (A-21245) (at 1:250 dilution), and anti-mouse AF488 at (A32723, 1:500 dilution) (all Thermo Fisher Scientific). Control experiments were performed by omitting primary antibodies from the immunostaining procedure.

Acid fuchsin orange G (AFOG) staining was performed using the AFOG staining kit (Biognost) following the manufacturer’s protocol.

For Phalloidin staining, 60 µm-thick tissue cryosections were air-dried, washed with PBS, and permeabilized for 2h in 0.5% Triton X-100/PBS. Sections were incubated with Phalloidin conjugated with AF-647 (A30107, Thermo Fisher Scientific) for 2h at a 1:1000 dilution (in 2% horse serum, 0.2% Triton X-100, and 1% DMSO in PBS). After 45 min wash with 0.1% Triton X-100/PBS, sections were mounted with fluorescence mounting medium (Dako, Agilent Technologies).

### Microscopy and image acquisition

Bright-field images were taken using ZEISS stereomicroscope Stemi 508 with a colour camera Axiocam 305 and Zen3.4 software. Images of whole heart sections were taken on fluorescent stereomicroscope Thunder Imager - model organism (Leica Microsystems) equipped with K8 CMOS camera and LAS X software. For higher magnifications, samples were imaged by Leica Confocal Microscope SP8 equipped with LAS AF software (Leica Microsystems). The acquisition of images was performed in sequential mode. Signal intensity was measured by the LAS AF quantification tool. Quantifications (scar resolution parameters, number of dedifferentiated CMs, length and number of protrusions) were done in LAS AF or Fiji (47) software.

### RNA isolation

Zebrafish ventricles from cryoinjured and sham-operated wt and *ankrd1a* mutant fish were homogenized in TRIzol Reagent (Invitrogen, Thermo Fisher Scientific) and total RNA was extracted using RNA Clean and Concentrator Kit (ZYMO Research), following the manufacturer instructions. Total RNA was eluted in 23 µl of DNase/RNase-free water and its quantity and purity were measured promptly after isolation, using BioSpec-nano (Shimadzu) and Qubit 4^TM^ Fluorometer with ssRNA BR Assay Kit^TM^ (Thermo Fisher Scientific). RNA samples were stored at -80°C for downstream use.

### Transcriptome analysis

Bulk mRNA-seq was performed using RNA isolated from the ventricles of cryoinjured and sham-operated wt and *ankrd1a* mutant zebrafish at 7 dpci. Three ventricles were pooled as one sample, and each group contained 4 samples. The quality of RNA was determined using the RNA 6000 Nano Kit (Agilent Technologies) and Bioanalyzer 2100 (Agilent Technologies). Samples with an RNA integrity number (RIN)≥ 5, concentration ≥ 20 ng/µl, and OD 260/280 between 1.7 and 2.1 were sequenced at Novogene (London, United Kingdom).

Messenger RNA was purified from total RNA via poly A enrichment. The cDNA libraries were prepared using the Illumina TruSeq Stranded mRNA Library Preparation kit (Illumina) and 150 paired-end sequencing was performed on the SEQ Novaseq 6000 platform (Illumina).

Raw reads of fastq format were processed using fastp (v0.23.4), applying the following parameters to remove residual adapter sequences and to keep only high-quality data (qualified_quality_phred=20, nqualified_percent_limit=30, average_qual=25, low_complexity_filter=True, complexity_threshold=30). Pared-end clean reads were mapped to the reference genome using STAR (v2.7.11b) with sjdbOverhang option set on read length and other parameters as default. Reference genome and transcripts’ annotation input files were downloaded from the Ensembl v112 repository. The percentage of uniquely mapped reads was high, with a mean value of 80,4%. To estimate transcript and gene abundances, alignments were analyzed by RSEM (v1.3.3), and the sample-specific gene-level abundances were merged into a single raw expression matrix by applying a dedicated RSEM command (rsem-generate-data-matrix). Before differential gene expression analysis, the raw expression matrix was normalized and filtered to include only genes with at least 10 counts in a number of samples (N), where N represents the sample size of the smallest experimental group by edgeR (v4.2.0) program package in R environment (v4.4.1). Exploratory data analysis was performed through box plot and principal component analysis (PCA). Differential gene expression analysis of two conditions was performed using a multi-factorial statistical analysis based on a negative binomial model that used a generalized linear model approach computed by edgeR (v4.2.0) from raw counts in each comparison (16 samples in 4 different groups). Benjamini & Hochberg method was used for multiple testing correction and genes with adjusted p values ≤ 0.05 and absolute fold change ≥ 0.5 were considered significantly differentially expressed. The bioMart package (v2.60.1) was used to annotate differentially expressed genes (DEGs), querying available Ensembl Gene IDs and retrieving Entrez gene IDs, Uniprot, and ZFIN gene symbols. The clusterProfiler (v.4.12.0) and org.Dr.eg.db as a reference database were used to perform DEG gene ontology enrichment analysis. Enriched terms with adjusted p values less than 0.05 were considered significantly enriched. Gene set pathway enrichment analysis was performed using ReactomePA (v 1.48.0). The results were visualized using R package gplots (v 3.1.3.1). *In silico* zebrafish matrisome list (48) was used to identify differentially expressed ECM proteins. Matrisome genes without Ensembl ID (68/1015) were not considered.

The RNA-seq data are deposited in NCBÍs Gene Expression Omnibus and are accessible through GEO Series accession number GSE285387, available at https://www.ncbi.nlm.nih.gov/geo/query/acc.cgi?acc=GSE285387.

### Real-time quantitative polymerase chain reaction (RT-qPCR)

Complementary DNA (cDNA) was synthesized using 0.5 µg of RNA and a High-Capacity cDNA Reverse Transcription Kit (Applied Biosystems), according to the manufacturer’s instructions. PCR reactions were performed in technical triplicate for each sample on Line-Gene 9600 Plus real-time PCR (Bioer Technology) using Power SYBR™ Green PCR Master Mix (Applied Biosystems). The rpl13a transcript was used as an internal reference for normalization (49). The data were analyzed by the −ΔΔCt method. Primers are listed in Table S1.

### Statistical analysis

Statistical analysis was conducted using GraphPad Prism software 6.0. The normal distribution and variance homogeneity of the data were verified by the Shapiro–Wilk and Levene’s tests, respectively. For individual comparisons between two independent experimental groups, the unpaired two-tailed Student’s t-test (for normally distributed data) or the Mann-Whitney test (for non-normally distributed data) was employed. Results are presented as mean ± SD, and a P-value ≤ 0.05 was considered significant. The Pearson correlation coefficient (r) was used to quantify the degree and direction of a relationship between two variables.

## RESULTS

### *TgBAC*(*ankrd1a:EGFP*) is expressed at 15 hpci in border zone CMs of cryoinjured zebrafish heart

In our previous work, we showed restricted *ankrd1a* expression in border zone CMs at 7 dpci (42). To determine the earliest time point of *ankrd1a* activation we assessed reporter fluorescence in injured hearts of *TgBAC(ankrd1a:EGFP);Tg*(-0.8*myl7*:*nls-DsRedExpress*) line with DsRed-labelled CMs nuclei. The EGFP was visible in the border zone from 15 hpci (Fig. 1).

**Figure 1.**
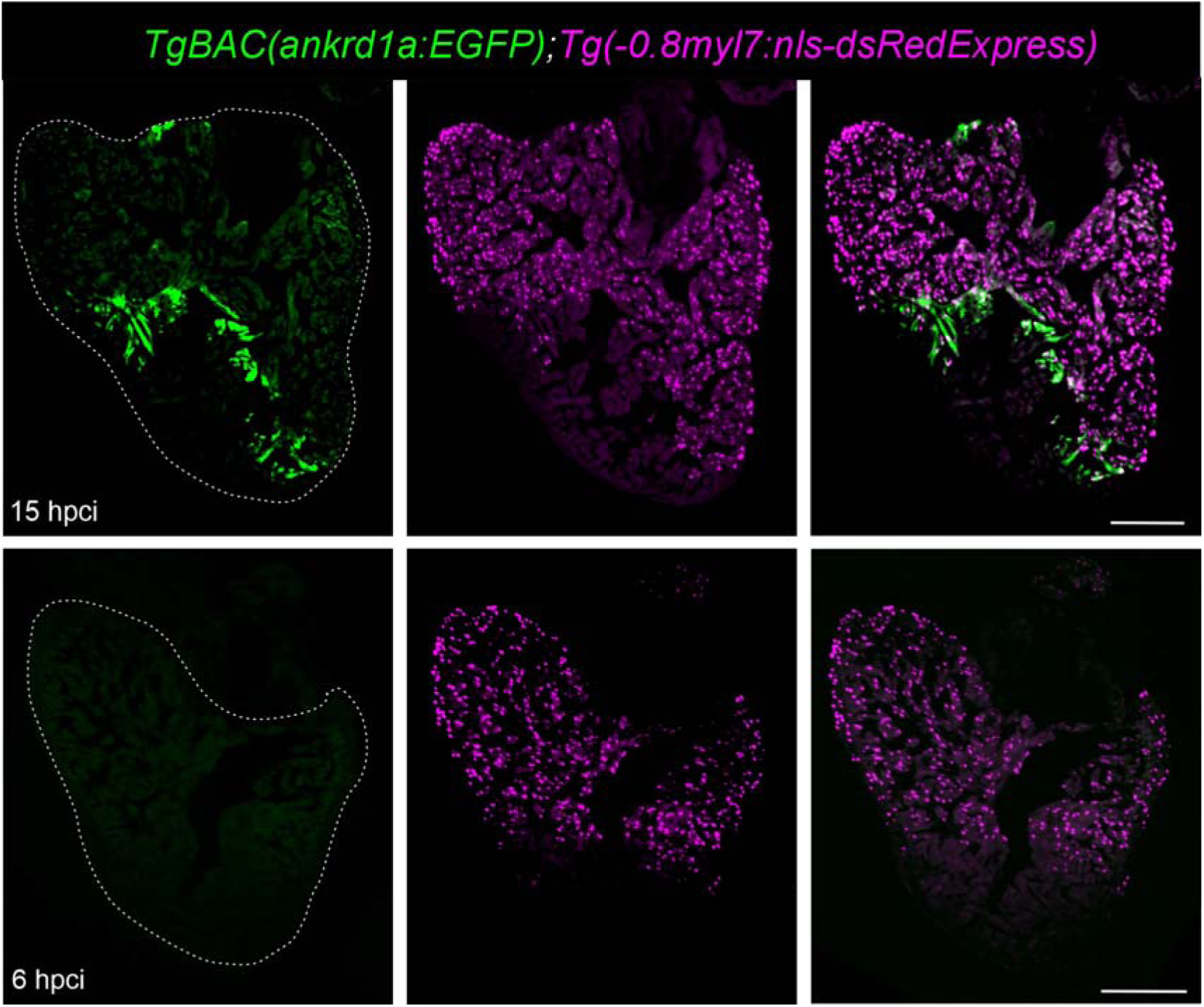
Early expression of *TgBAC*(*ankrd1a:EGFP*) transgene in the border zone CMs. Hearts of *TgBAC(ankrd1a:EGFP);Tg*(-0.8*myl7*:*nls-DsRedExpress*) (n=4) were crioinjured and left to recover for 6 and 15 h. White dashed line outlines the ventricle. Scale bars, 200 µm.

### *TgBAC*(*ankrd1a:EGFP*) is not expressed in the epicardium and endocardium of cryoinjured zebrafish heart

Since *ankrd1a* expression was not detected in cardiac cells other than CMs at 7 dpci, we explored its expression in endocardium and epicardium at 3 and 10 dpci. The activation of *ankrd1a* in the epicardium of the regenerating heart was tested by Caveolin-1 staining to identify epicardial cells in hearts of *TgBAC(ankrd1a:EGFP)* line. The EGFP signal was visible in the injury border zone, but there was no overlap with the Caveolin-1 signal (Fig. 2A).

**Figure 2.**
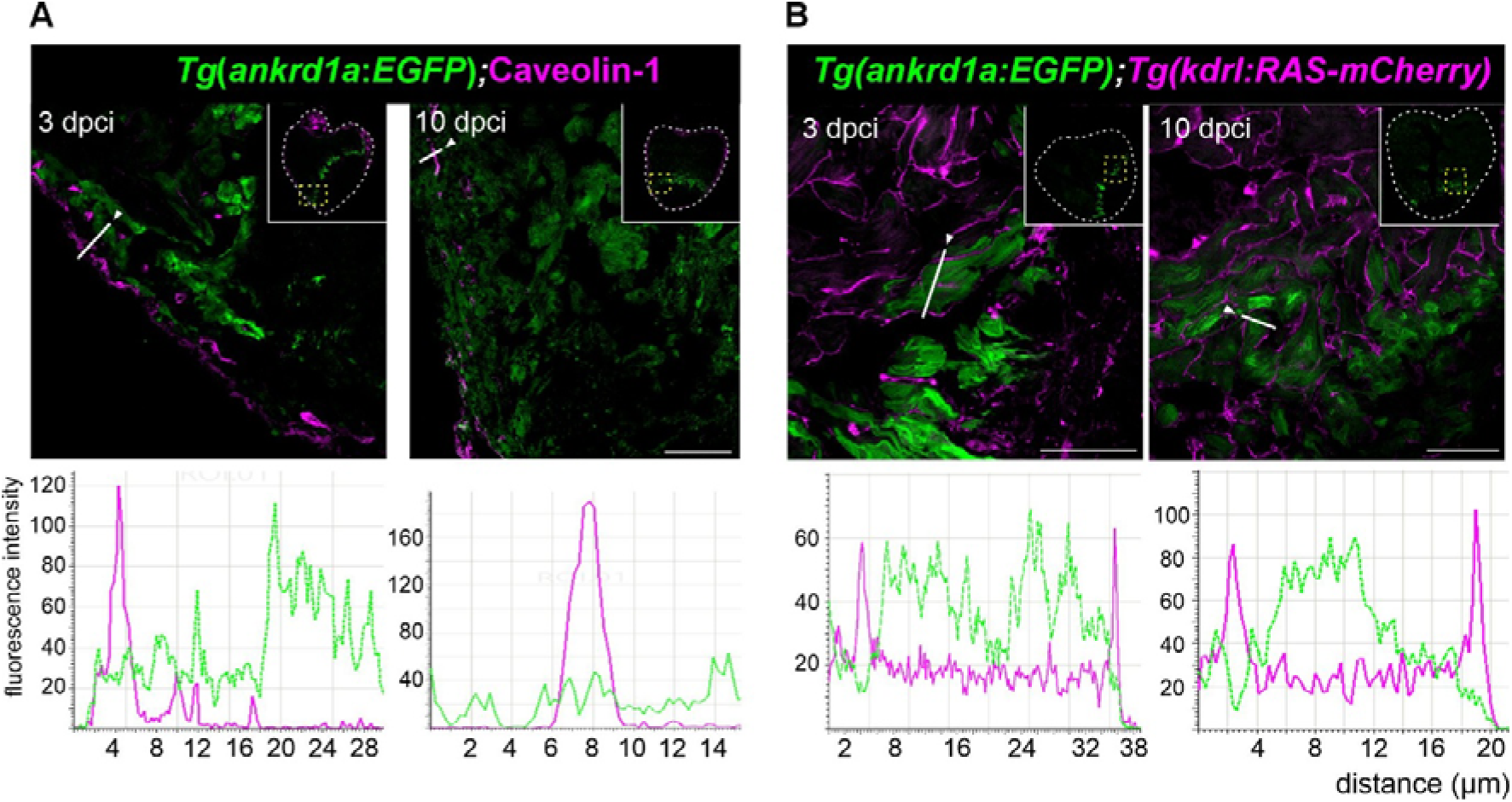
There is no *TgBAC*(*ankrd1a:EGFP*) expression in the epicardium and endocardium of the regenerating zebrafish heart. Hearts (n=3 per time point) of (A) *TgBAC(ankrd1a:EGFP)* and (B) *TgBAC(ankrd1a:EGFP);Tg(kdrl:RAS-mCherry)* were cryoinjured and analysed 3 and 10 days after cryoinjury. (A) Cryosections were stained for EGFP (green) and Caveolin-1 (magenta). (B) Representative images of cryosection showing EGFP positive (green) and mCherry positive (magenta) cells. White dashed lines in insets outline the ventricles. Yellow dashed rectangles in insets show the position of high magnification images. Bottom panels: fluorescence intensity (in arbitrary units) profiles of ankrd1a:EGFP (green line) and Caveolin-1 (A) or kdrl:RAS-mCherry (B) (magenta line) measured along the selected region (white lines) shown on the top images. Scale bars, 50 µm for A and 25 µm for B.

To investigate if *ankrd1a* gets activated in the endocardium of injured heart, we employed the *Tg(kdrl:RAS-mCherry)* reporter line in which membranes of endocardial cells are labelled with a fluorescent marker. In double reporter line *TgBAC(ankrd1a:EGFP)*;*Tg(kdrl:RAS-mCherry)*, at 3 and 10 dpci, there was no EGFP and mCherry signal colocalization (Fig. 2B).

### *TgBAC*(*ankrd1a:EGFP*) is expressed in dedifferentiating and proliferating CMs in cryoinjured heart

Since *ankrd1a* upregulation in border zone CMs was previously shown (42), we investigated whether these *ankrd1a* positive CMs undergo dedifferentiation and proliferation, the hallmark processes occurring in the border zone during regeneration of zebrafish heart. Heart cryosections of *TgBAC*(*ankrd1a*:*EGFP*) were stained for EGFP and embryonic cardiac myosin heavy chain (embCMHC) at 1 and 7 dpci. Colocalization of embCMHC and EGFP was observed at 7, while at 1 dpci the transgene expression preceded that of dedifferentiation marker embCMHC (Fig. 3).

**Figure 3.**
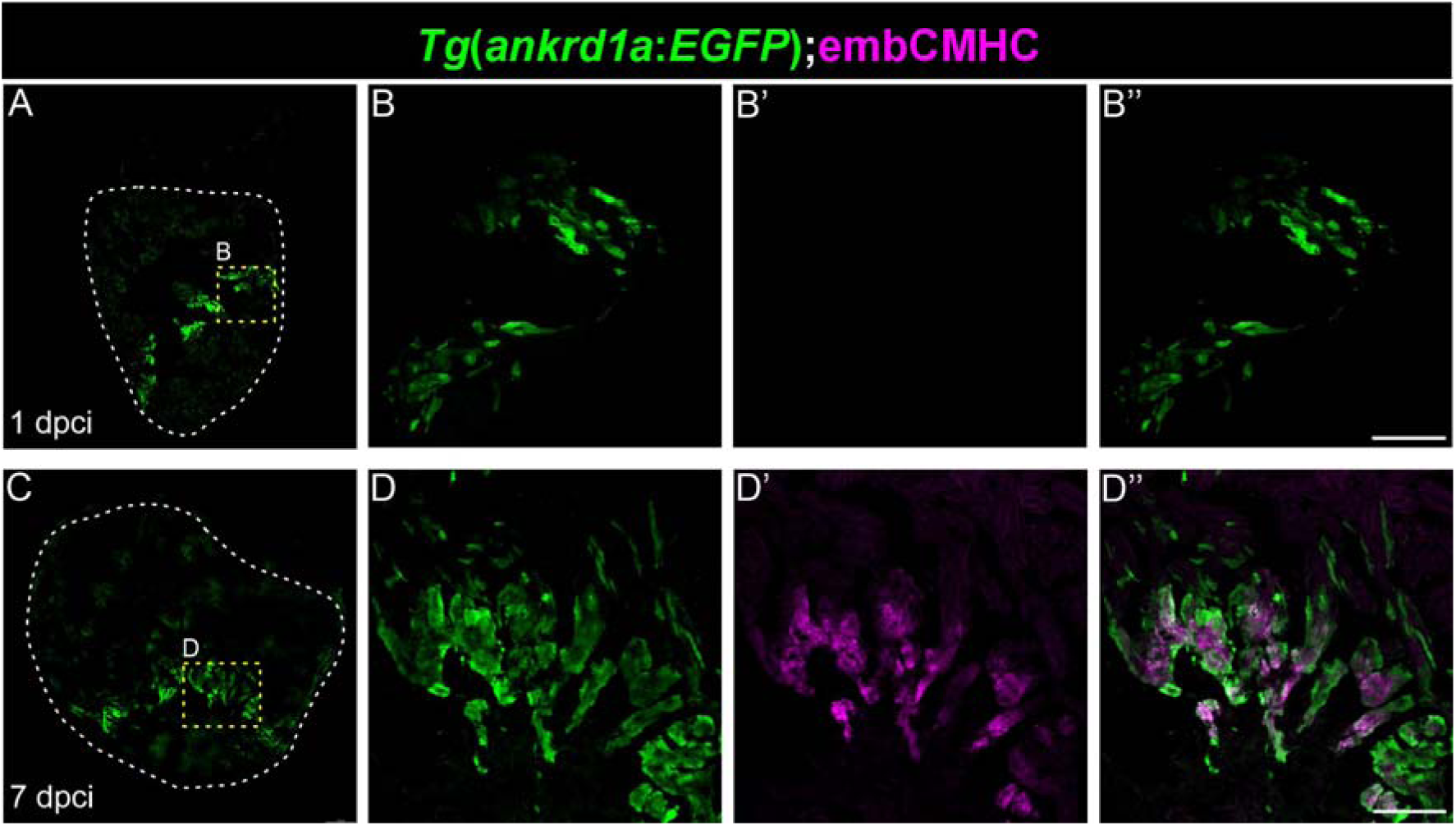
TgBAC(ankrd1a:EGFP) is expressed in dedifferentiating CMs upon heart cryoinjury. Cryosections of injured hearts (n=3 per time point) were stained for EGFP (green) and embCMHC (magenta). White dashed lines on A and C outline the ventricle. Insets B and D (yellow dashed rectangles) show the position of high magnification images (B-B’’ and D-D’’) of the border zone CMs. Scale bar, 50 µm.

To test whether *ankrd1a-*positive CMs proliferate, heart cryosections of double reporter line *TgBAC(ankrd1a:EGFP);Tg(myl7:nls-DsRedExpress)* were stained for cell proliferation marker Pcna. At 7 dpci, when the proliferation of CMs peaks, we observed EGFP^+^/DsRed^+^/Pcna^+^ proliferating CMs expressing *ankrd1a* (Fig. 4). However, not all EGFP-positive CMs had proliferative status (EGFP^+^/DsRed^+^/Pcna^-^), and there were proliferative CMs not expressing ankrd1a:EGFP transgene (EGFP^-^/DsRed^+^/Pcna^+^) at 7 dpci (Fig. S1), suggesting that *ankrd1a* activation is not restricted to proliferating CMs.

**Figure 4.**
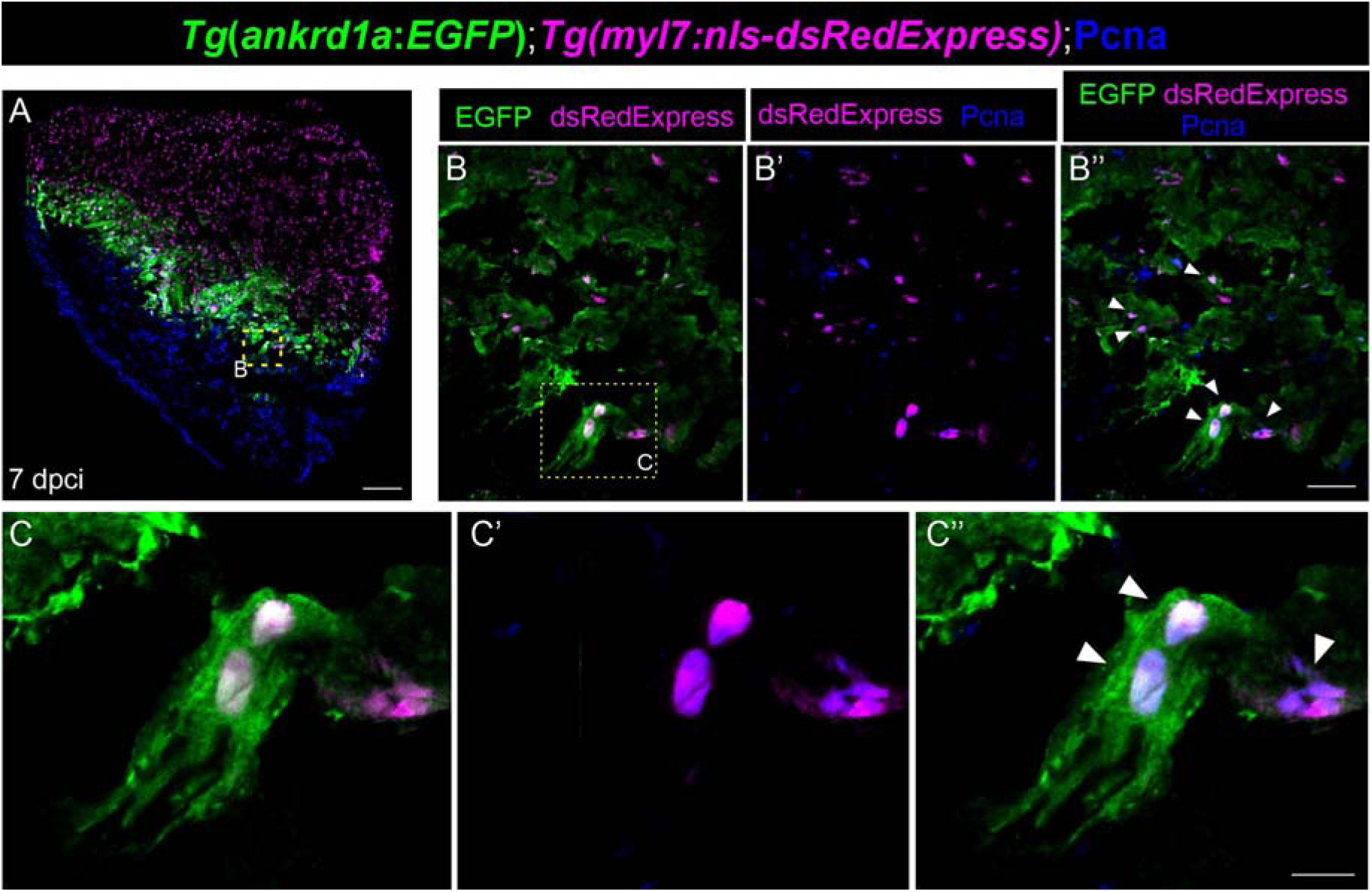
***TgBAC(ankrd1a:EGFP)* is expressed in proliferating cardiomyocytes**. A) Hearts (n=3) of *TgBAC(ankrd1a:EGFP);Tg(myl7:nls-DsRedExpress)* were injured and left to recover for 7 days. Cryosections were stained for EGFP (green), DsRed (magenta) and proliferation marker Pcna (blue). Insets (yellow dashed lines) on A and B show the position of high magnification images on B-B’’ and C-C’’, respectively. White arrowheads point to proliferating cardiomyocyte nuclei. Scale bars, 100 µm for A, 25 µm for B, and 10 µm for C.

### *TgBAC*(*ankrd1a:EGFP*) is expressed in invading CMs with protrusions

To replenish lost myocardium, newly proliferated CMs migrate and invade collagen-containing fibrotic scar tissue. Using double transgenic line *TgBAC*(*ankrd1a*:*EGFP*);*Tg*(*myl7*:*nls-DsRedExpress*) we detected EGFP in CMs with protrusions at 10 dpci (Fig. 5), when their number and length reach a peak.

**Figure 5.**
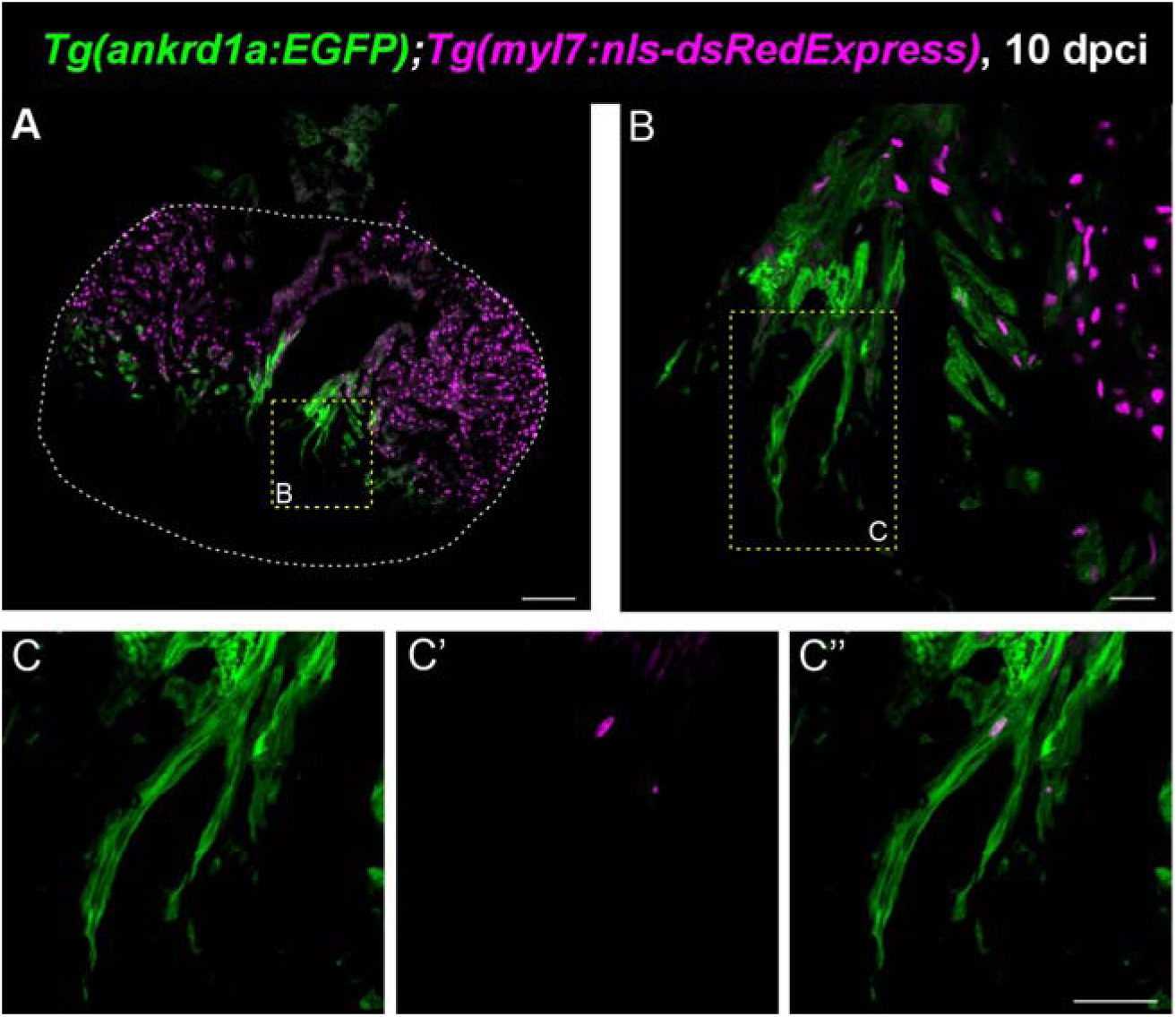
***TgBAC(ankrd1a:EGFP)* expression in the CMs with protrusions of the regenerating heart.** A) Sagittal section of injured *Tg(ankrd1a:EGFP);Tg(myl7:nls-DsRedExpress)* adult heart (n=3) left to recover 10 days after cryoinjury. The white dashed line outlines the ventricle. Insets B and C (yellow dashed lines) show the position of high magnification images (B and C-C’’, respectively) of CMs with (most prominent) protrusions. Scale bars, 100 µm for A, 20 µm for B, and 25 µm for C-C’’.

### *TgBAC*(*ankrd1a:EGFP*) is expressed in cells lining the residual scar

To assess *ankrd1a* activation at later time points after cryoinjury, we analysed hearts of *TgBAC(ankrd1a:EGFP);Tg(-0.8myl7:nls-DsRedExpress)* zebrafish at 30 dpci. In all hearts with incomplete scar resolution, EGFP-positive CMs were bordering the collagenous scar independently of the remaining injury size (Fig. 6A-D). The intensity of the EGFP signal was visibly higher in zebrafish with an extensive scar and low regeneration progression (Fig. 6A-A’’) and partially regenerated myocardium (Fig. 6B-B’’). In hearts where the regeneration has progressed more (Fig. 6C-C’’) or was almost complete (Fig. 6D-D’’), the EGFP signal was of lower intensity, but still localized along residual traces of collagen. No expression of *ankrd1a:EGFP* transgene was observed in the CMs of the regenerated ventricular tissue, which is visible as a thickening of the cortical layer of the myocardium.

**Figure 6.**
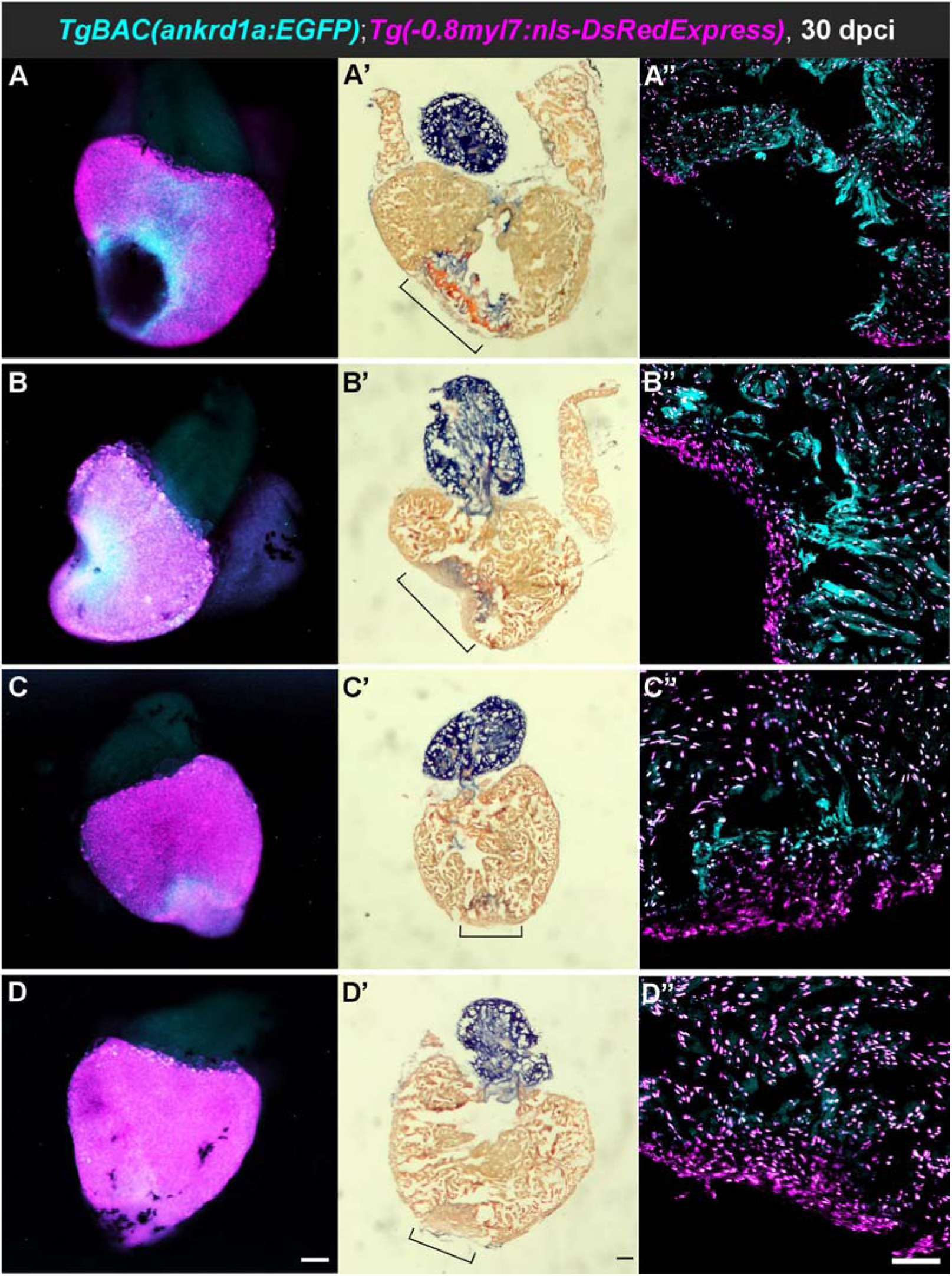
***TgBAC*(*ankrd1a:EGFP*) expression is related to the residual scar size.** Hearts (n=4) of *TgBAC(ankrd1a:EGFP)*;*Tg(-0.8myl7:nls-DsRedExpress)* fish were cryoinjured and analyzed at 30 dpci, showing different levels of scar resolution (least (A) to most (D) regenerated). The whole hearts after dissection were imaged by fluorescence microscopy (A-D). Serial sections were used for AFOG staining (bright field images on panels А’-D’) and fluorescence imaging by confocal microscopy (А’’-D’’). In the indicated region of injury on panels A’-D’, collagen is stained blue, fibrin is stained orange, and CMs are stained brown. Scale bars, 100 μm.

### *ankrd1a* mutant regenerates cryoinjured hearts

Recently, we reported the generation of *ankrd1a* mutant zebrafish line, which was used to investigate the role of this gene in skeletal muscle repair after needle stab injury (45). To analyse whether the loss of *ankrd1a* function affects the outcome of cardiac regeneration, we monitored the scar resolution process in injured *ankrd1a* mutant and wt zebrafish hearts. Histological sections were stained with AFOG at 30 dpci, to distinguish cells, collagen, and fibrin. Light microscopy images were used to quantify three scar resolution-related parameters in the heart sections: ventricle area, collagen deposit area, and regenerated tissue area. From these measurements, the following values were calculated: 1) total injury area as a sum of collagen deposit area and regenerated tissue area, 2) relative scar area as a ratio between collagen deposit area and ventricle area, and 3) relative regenerated area as a ratio of regenerated tissue area and total injury area (Table S2). There was no statistically significant difference for all six analysed parameters between wt and *ankrd1a* mutant zebrafish, suggesting that loss of *ankrd1a* function does not affect scar resolution during regeneration of zebrafish heart (Fig. 7A). Representative heart sections show neither structural differences in the regenerated myocardial regions (Figure 7B, left panels), nor in the form of residual scarring (Figure 7B, right panels) between wt and *ankrd1a* mutant zebrafish.

**Figure 7.**
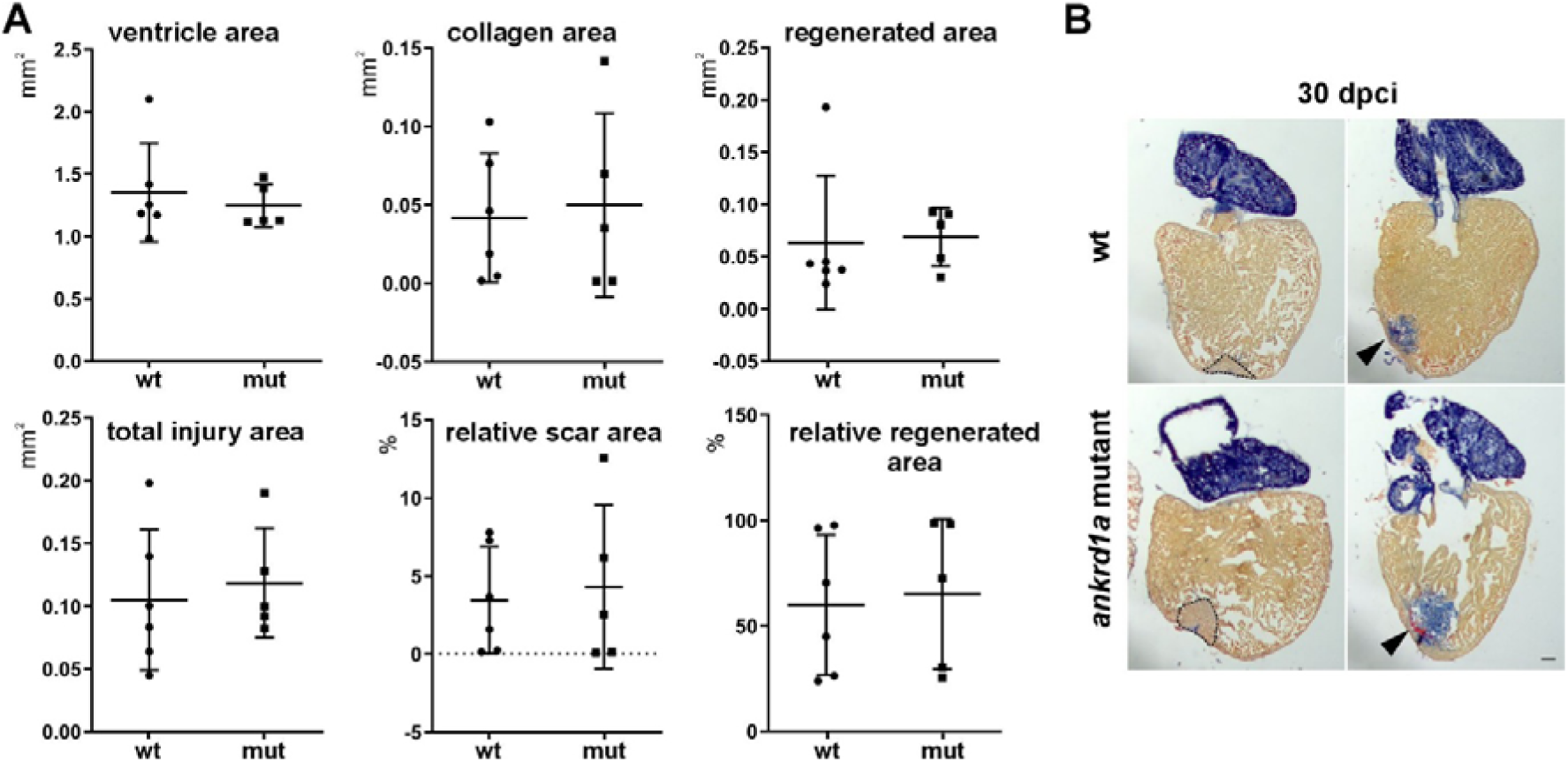
Scar resolution during zebrafish heart regeneration is not affected by the loss of *ankrd1a* function. A) Measured values for ventricle area, collagen deposit area, and regenerated tissue area, and calculated values for total injury area, relative scar area, and relative regenerated area were compared between the *ankrd1a* mutant (n=5) and wt (n=5) hearts at 30 dpci. P>0.1 for all comparisons. B) Representative images of wt and ankrd1a mutant heart cryosections stained with AFOG kit. Collagen is blue, fibrin is orange, and CMs are brown. Black dashed lines outline areas of regenerated myocardial tissue, arrowheads point to unresolved scars. Scale bar, 100 μm.

### Immune response and cell adhesion are affected in the regenerating hearts of the ankrd1a mutant

Since *ankrd1a*, although not essential, finely regulates skeletal muscle repair (45), we performed transcriptome profiling to identify targets of *ankrd1a* involved in heart regeneration. RNAseq was done for cryoinjured ventricles of *ankrd1a* mutant and wt zebrafish at 7 dpci, while hearts of sham-operated fish of both genotypes served as controls. To visualize the major changes in gene expression, we performed principal component analysis (PCA) based on normalized counts of all genes expressed in the hearts of cryoinjured and sham-operated fish of both genotypes. PCA showed that samples were separated according to experimental conditions in component 1 (∼30% of variability), and genotype in component 2 (∼14.5% of variability) (Fig 8A). Unique and shared DEGs between different conditions are presented as a Venn diagram (Fig. 8B).

**Figure 8.**
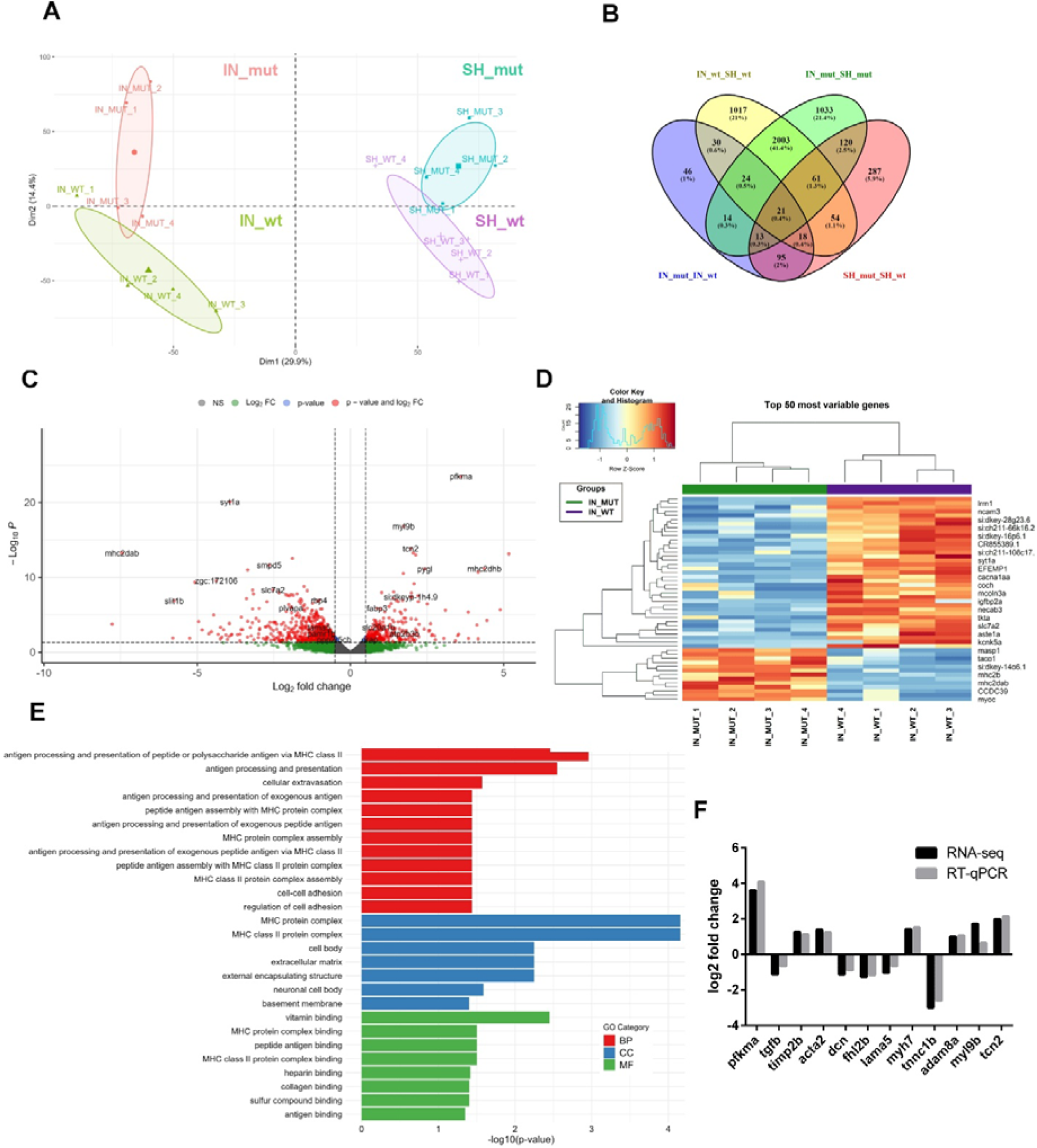
Transcriptome analysis of cryoinjured hearts of *ankrd1a* mutant and wt zebrafish at 7 dpci. A) PCA performed on the normalized counts of sham-operated (SH) and cryoinjured (IN) *ankrd1a* mutant (mut) and wt (wt) hearts (n=4 for each experimental group, 3 pooled ventricles in each sample). The first two dimensions capture ∼45% of the variability of the data; B) Venn diagram indicating the number of overlapping DEGs among all four groups. The numbers of genes in each intersection are indicated; C) Volcano plot showing DEGs between cryoinjured *ankrd1a* mutant and wt hearts, red dots – significantly differentially expressed genes, blue dots – significant genes based on the p value, green dots – significant genes based on the log2 fold change, grey dots – non-significant genes; D) Heatmap of the top 50 DEGs between cryoinjured heart of *ankrd1a* mutant and wt showing expression patterns across samples. The Z-scores of gene expression measurements are displayed as colours ranging from blue to red; E) Summary of the most enriched gene ontology terms in cryoinjured *ankrd1a* mutant versus wt hearts by category. BP - biological process, CC - cellular compartment and MF - molecular function; F) Comparison of expression levels of 12 selected DEGs measured by RNA-seq and RT-qPCR. Results are presented as log2 fold change.

The differential gene expression (DEG) analysis identified 261 genes that were significantly upregulated (177) or downregulated (84) in the *ankrd1a* mutant compared to wt at 7 dpci (Fig. 8C and D, Table S3). Gene ontology analysis showed a notable enrichment of genes associated with antigen processing and presentation (*mhc2dab,mhc2dhb,mhc2dgb,mhc2daa,mhc2dga*), cellular extravasation (*plvapb, plvapa, adam8a*), and cell-cell adhesion (*itgb1a*, *itgam.1, pcdh1g18, pcdh1g2, ncam3, cxadr, lama5, ptprfb, adam8a, pcdh15a*) (Fig. 8E, Table S4). *In silico* analysis of the zebrafish matrisome identified 26 DEGs, comprising 13 core matrisome genes, 12 matrisome-associated genes, and 1 putative matrisome gene (Table S5). Among them, 11 genes encode ECM glycoproteins, 5 encode ECM regulators and 4 encode secreted factors.

Technical validation of transcriptomic results was performed by RT-qPCR for 12 protein-coding genes. They are muscle-specific (*myh7*, *tnnc1b*, *myl9b*, *acta2*, *fhl2b*), related to ECM (*timp2b*, *dcn*, *lama5*, *adam8a*), metabolism (*pfkma*), cell division (*tgfb5*), and cellular uptake of cobalamin (*tcn2*). The qRT-PCR results were in line with the RNA-seq data (Fig. 8F), with a Pearson correlation coefficient r=0.974, confirming the reliability of the transcriptomic analysis.

There were 3228 genes differentially expressed between cryoinjured and sham-operated hearts of wt (3026 up- and 202 down-regulated) (Fig. S2A, Table S3) and 3289 genes differentially expressed between cryoinjured and sham-operated hearts of *ankrd1a* mutant (2970 up- and 319 down-regulated) (Fig. S2B, Table S3). Those DEGs are involved in actin filament organization, mitotic cell cycle process, regeneration, extracellular matrix organization and other processes important for heart regeneration (Table S4). When compared, there is significant overlap (2109 genes) between these two lists of DEGs, indicating that heart regeneration is not critically affected by the *ankrd1a* loss of function.

Comparison of transcriptomes between sham-operated wt and *ankrd1a* mutant showed 669 DEGs (267 up- and 402 down-regulated) (Fig. S2C, Table S3). Those genes are not enriched in any biological process based on gene ontology analysis, but their molecular function is related to vitamin binding, MHC protein complex binding and protein phosphatase activity (Fig. S2C, Table S4).

### Loss of the *ankrd1a* function affects the number of dedifferentiating CMs

Since *myh7* (ENSDARG00000079564, previously *vmhc*), known to be upregulated in dedifferentiating border zone CMs (13), was identified as one of the upregulated DEGs in the *ankrd1a* mutant (FC=2.67), we tested whether loss of *ankrd1a* function affects CMs’ dedifferentiation. Cryoinjured hearts were immunostained with the N2.261 antibody (specific to embCMHC) to mark dedifferentiated (and undifferentiated) cardiomyocytes in *ankrd1a* mutant and wt, at 7 dpci (Fig. 9). Compared to wt, there were significantly more N2.261^+^ CMs in the border zone of *ankrd1a* mutant hearts (24.327±4.818% for wt and 32.103±5.816 for mutant), suggesting increased CMs dedifferentiation.

**Figure 9.**
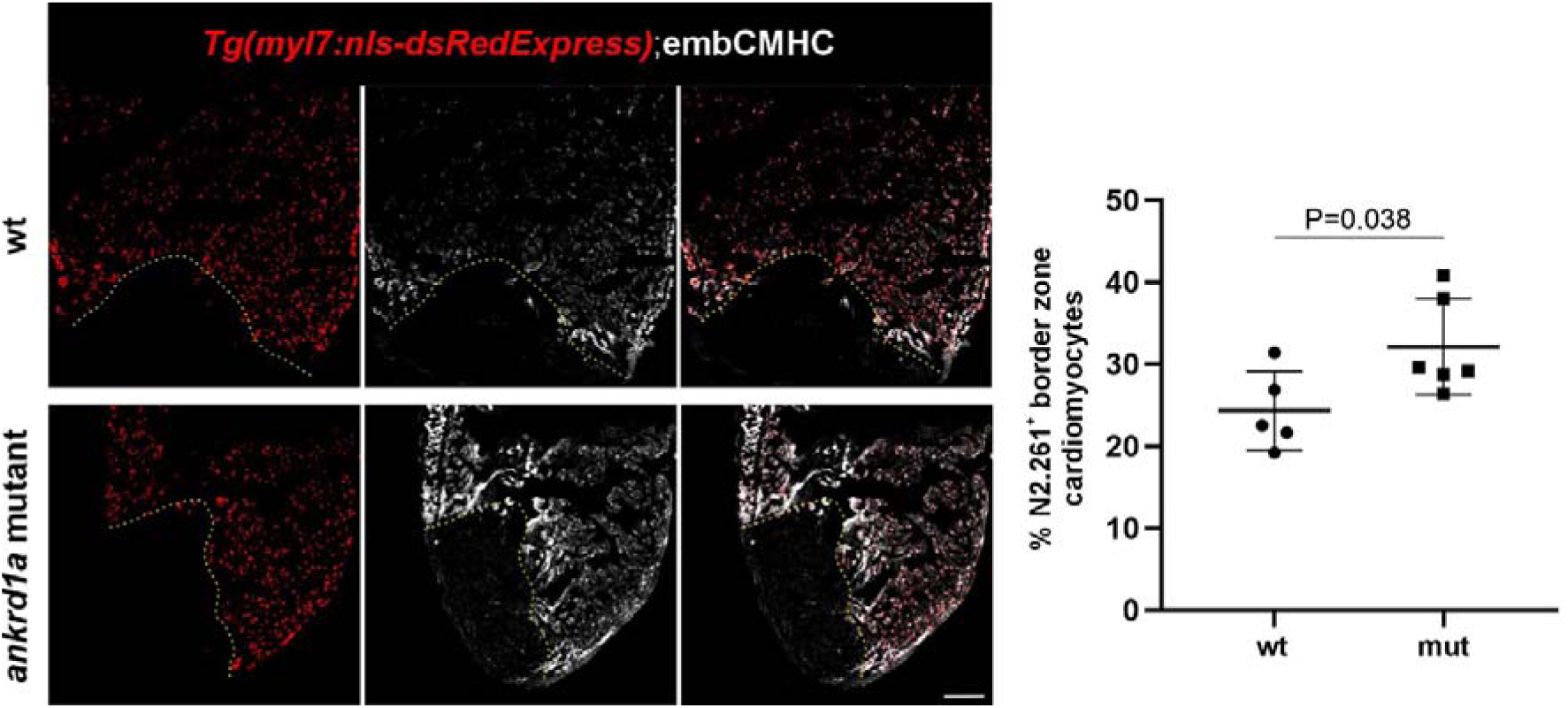
The number of dedifferentiating CMs is increased in the injured *ankrd1a* mutant heart. Images of representative cryosections of wt (n=5) and *ankrd1a* mutant (n=6) zebrafish ventricles at 7 dpci, immunostained for DsRed and N2.261, showing dedifferentiating cardiomyocytes (left), and respective quantification (right). Cortical CMs were excluded from the analysis. Quantification of CMs dedifferentiation was performed in the area 100 μm proximal from the injury border in three non-consecutive sections with the largest injured area from each heart. Yellow dashed lines mark the border of the injured tissue. Dots in the graphs represent individual ventricles. Results are presented as meanL±LSD. P-value is calculated using unpaired two-tailed Student’s t-test. Scale bar, 100 μm.

### Loss of *ankrd1a* expression does not affect the number and length of CMs protrusions

Transcriptome analysis revealed altered expression of ECM-related genes in *ankrd1a* mutant heart at 7 dpci. Since ECM remodelling is associated with invasion of CMs into injured tissue, we tested whether the loss of *ankrd1a* affects the number and length of protrusions in these repopulating CMs. Thick (60 μm) heart cryosections obtained from the *ankrd1a* mutant and wt zebrafish at 10 dpci were stained with Phalloidin, to detect F-actin (Fig 10A), and protrusion lenght and number were quantified (Fig.10B and C, respectively) as recently reported (50). There was no significant difference either in the average length of protrusions (25.95±4.93 µm for wt and 31.68±5.68 µm for *ankrd1a* mutant) or in their number per 100 µm of the border zone (3.76±0.54 for wt and 3.59±0.70 for *ankrd1a* mutant), indicating no perturbation in CMs invasion caused by the loss of *ankrd1a* function.

**Figure 10.**
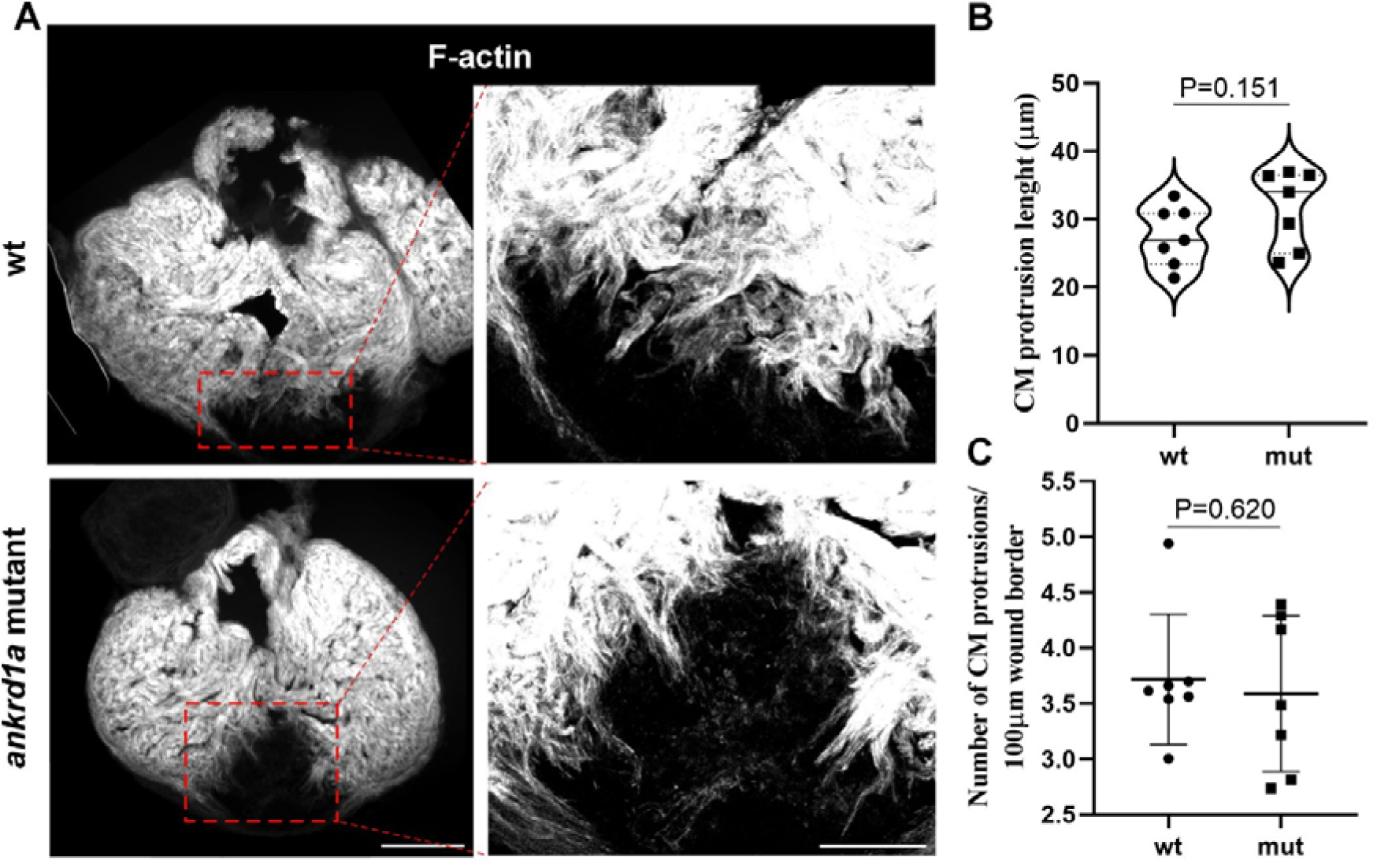
The number and length of CM protrusions are not influenced by the loss of *ankrd1a* function. (A) Thick cryosections of *ankrd1a* mutant and wt zebrafish ventricles (n = 7 for both wt and *ankrd1a* mutant) at 10 dpci were stained with Phalloidin to detect F-actin. CM protrusions were quantified using a maximum intensity projection image from at least 2 sections showing the largest injured area. Protrusions of trabecular CMs were used for quantification. The red dotted box denotes the zoomed image from the wound border zone. Scale bars, 250 μm for left (lower magnification images) and 100 μm for right (higher magnification images) panels. Quantification was done for the (B) average length of CM protrusions per 100 μm of wound border and (C) their number. For the length of CM protrusions, data are presented as violin plots, with solid lines indicating the median and dotted lines indicating the 25^th^ and 75^th^ percentiles. For the number of CM protrusions, data are presented as meanL±LSD. Dots in the graphs represent individual ventricles. P-values were calculated using unpaired two-tailed Student’s t-test for the length and Mann-Whitney test for the number of protrusions.

## DISCUSSION

In our previous work, using *TgBAC(ankrd1a:EGFP)* reporter line (42), we showed upregulation of *ankrd1a* in border zone CMs, where post-injury tissue remodelling occurs (42). In human and mouse infarcted hearts, ANKRD1 also marks the injury border zone, which is transcriptionally distinct and spatially confined from the remote myocardium (51). To make a step toward deciphering *ankrd1a* function in zebrafish heart regeneration, we explored its activation in different cardiac structures, at different time points after cryoinjury and identified targets of this gene in the regenerating heart by transcriptome profiling.

We found *ankrd1a* activated at 15 hpci, which makes him one of the earliest markers of activated border zone CMs in zebrafish heart after cryoinjury. Its early activation was also shown in a murine model of myocardial infarction (MI), where Ankrd1 was found as one of the mechanical stress-response genes highly expressed in a unique cluster of border zone CMs at one day post MI. Integrative analysis of snRNA-seq and spatial transcriptome suggested preventive role of mechanosensing genes upregulated in the border zone during the acute phase after MI against post-MI left ventricular remodelling (52). It remains to elucidate which factors drive upregulation of these early-response mechanosensitive genes, and whether *ankrd1a* has a specific role in heart muscle remodelling, as demonstrated for *Csrp3* (52). It is important to note that zebrafish homolog *csrp3* and a few other mechanosensitive genes (*flnc*, *xirp1*, *fhl2*) are upregulated only in the *ankrd1a* mutant (but not in wt) heart after the cryoinjury at 7 dpci.

The epicardium and the endocardium contribute to the regenerative response to cardiac injury in the zebrafish by providing mechanical and paracrine support, and they are activated before the myocardium and the coronary vasculature (53–56). Here we tested the *ankrd1a* expression in epicardium and endocardium, showing that these cardiac layers can be excluded as the sites of *ankrd1a* expression at the region of injury. Transcriptomics dataset from Weinberger et al (57), informing that *ankrd1a* is not differentially expressed in the epicardium of the injured heart, corroborates our findings.

Predominant expression of *ankrd1a* in CMs of the border zone imposed the question of its involvement in dedifferentiation, proliferation and migration of CMs, key events in regenerating zebrafish heart, leading to replenishment of lost cardiac cells. Although *ankrd1a* activation was observed in CMs expressing markers of dedifferentiation (embCMHC) and proliferation (Pcna), the transgene was also found in embCMHC-negative and Pcna-negative CMs of the border zone, suggesting that *ankrd1a* expression is not exclusively coupled with these two processes. However, even at a later stage of regeneration (10 dpci), *ankrd1a* activation was still associated with the border zone, namely with a population of newly proliferated CMs with protrusions that migrate and invade the fibrotic scar tissue, enabling structural and functional replenishment of the injured tissue.

We further examined expression of *ankrd1a* at 30 dpci, when scar tissue removal is still ongoing and residual scars of different sizes are present (6, 23, 58–60). In hearts with incomplete scar resolution, we identified a distinct population of CMs expressing *ankrd1a*, consistently bordering residual collagen and fibrin. The transgene expression level was related to the delay in regeneration, manifested by the residual scar size. It appears that injury border zone CMs maintain their specific expression profile, at least regarding *ankrd1a*, during the scar removal process, and that *ankrd1a* expression at the injury site marks incompletely recovered cardiac tissue. The expression of *ankrd1a* in CMs at the border with non-contractile tissue, at an advanced stage of regeneration may be a response to the altered mechanical forces to which these cells are exposed due to the disturbed structure and geometry of the ventricular wall (61). This hypothesis is supported by our results of the forced swimming experiment (42), showing non-uniform upregulation of *ankrd1a* in ventricular CMs after increased loading of the heart muscle. It might also explain *ankrd1a* expression in the border zone within 15 hpci, as a response to abrupt change in ventricular muscle mechanics. Additionally, it was proposed that ANKRD1 regulates passive force by locking titin to the thin filament and that stressed muscle uses this mechanism to protect the sarcomere from mechanical damage in humans (27, 28).

From our comprehensive expression profiling, we can conclude that *ankrd1a* activation occurs throughout all phases of zebrafish heart regeneration, from border zone formation to scar resolution, characterising it as a consistent marker of CMs bordering injured tissue.

Regarding the function of *ankrd1a*, we showed that this gene is not essential for cardiac regeneration; injured hearts of the *ankrd1a* mutant regenerated similarly to wt. We recently also revealed its redundancy in skeletal muscle repair (45). Despite the lack of obvious phenotype in the *ankrd1a* mutant and its ability to regenerate injured striated muscle, we were interested in identifying potential changes in molecular processes caused by the loss of *ankrd1a* function during cardiac regeneration. Transcriptome analysis was performed at 7 dpci, when the most prominent events at the border zone are CMs dedifferentiation and proliferation. Among 261 DEGs, we found myosin *myh7* significantly upregulated in the injured *ankrd1a* mutant compared to wt. Dedifferentiating cardiomyocytes reactivate embryonic myosin isoforms, particularly *myh7*, previously designated as *vmhc* (ventricular myosin heavy chain orthologous to several human genes including *MYH7*) in regenerating zebrafish hearts (13). We showed that the number of dedifferentiating CMs in the *ankrd1a* mutant was increased, suggesting the potential role of *ankrd1a* in the regulation of CMs dedifferentiation in regenerating zebrafish heart. The role of *ankrd1a* in myogenic differentiation (62) has been recently corroborated by our finding that differentiation of newly forming myofibers in zebrafish was accelerated in *ankrd1a* mutant (45). It is possible that *ankrd1a* also fine-tunes cardiac regeneration by preventing excessive dedifferentiation of CMs.

Transcriptome analysis showed no changes in expression of cell cycle and proliferation-associated genes in *ankrd1a* mutant compared to wt fish. Recently, Liu et al (63) reported the regulatory role of Ankrd1 in cardiac regeneration in mice by modulating CMs proliferation. Our results in zebrafish are not in line with these findings. It is unclear whether this discrepancy is due to differences between injury models used in these two studies (cryoinjury and apical resection) or it reflects the differences in mechanisms of cell cycle control in CMs of adult fish and neonatal and adult mice.

According to transcriptome analysis the major biological processes affected by the loss of *ankrd1a* function at 7 dpci were related to antigen processing and presentation of peptide or polysaccharide antigen via MHC class II, as well as cell adhesion. It is well documented that immune response is essential for zebrafish heart regeneration (64–66). In particular, antigen-presenting cells (APCs) such as dendritic cells and macrophages play central roles in modulating this process by presenting antigens to adaptive immune cells and shaping the local inflammatory environment to support tissue repair. Their activity contributes to both immune resolution and CMs proliferation (64, 65). Dendritic cells are predominant APCs in cardiac tissue (67), while macrophages influence CM behaviour by secreting cytokines that regulate proliferation, hypertrophy, and electrical conduction. Conversely, CMs can secrete cytokines to recruit macrophages and modulate their phagocytic activity (68). Evidence also suggests close physical interactions between macrophages and protruding border zone CMs in the injured heart, where macrophages contribute to ECM remodelling (50). When MHC class II antigen presentation was disrupted in injured zebrafish heart, T cells and activated endocardial cells failed to efficiently populate the injury site, reducing CMs dedifferentiation and proliferation (69).

So far, there is no experimental evidence that the function of ANKRD1 is related to immune response. However, *Ankrd1* was identified as an immune-related gene in the study which used animal experiments and bioinformatics analysis to identify left ventricular hypertrophy (LVH)-related immune genes (70). *Ankrd1* and other hub genes were strongly correlated with fibroblasts and macrophages in immune analysis and positively associated with LVH in the experimental validation.

The question is in which cell type *ankrd1a* regulates the expression of genes related to antigen presentation. One possibility is CMs themselves, as the major cell population in regenerating heart expressing *ankrd1a*. Although CMs are not typically considered as antigen-presenting cells, they can express MHC class II genes upon induction, such as cardiac injury and regeneration (71). On the other hand, it is also possible that *ankrd1a* might be expressed in immune cells and exert a role in immune response during cardiac regeneration. In a study aimed at discovering the composition and activation states of immune cell populations of the regenerating heart (manuscript in preparation by Filipa Simões group), single-cell transcriptomic analysis at 5 dpci showed *ankrd1a* expression in plasmacytoid dendritic cells (pDCs), a subset of dendritic cells with a potential role in cardiovascular diseases (72–74). Although our investigation of *ankrd1a* expression did not include APCs or other immune cells, future studies should focus on deciphering the role of *ankrd1a* in immune response during heart regeneration.

There were 26 genes related to ECM among DEGs identified in cryoinjured *ankrd1a* mutant hearts (compared to wt) by transcriptome analysis. This prompted us to test whether the loss of *ankrd1a* function might affect CMs with protrusions that express ECM structural and remodelling proteins (9). Neither the length nor the number of CMs protrusions at 10 dpci were significantly affected in the *ankrd1a* mutant. However, the possible role of *ankrd1a* in ECM remodelling could not be excluded since ANKRD1 was found to regulate the expression of matrix metalloproteinase MMP13 (75) and motility of skin fibroblasts through interaction with a collagenous matrix (76).

Taken all together, presented results are in line with the hypothesis of the *ankrd1a* as a stress sensor (11, 42, 45). A “loss-of-neighbour” hypothesis (77) proposes that non-injured CMs at the border zone are mechanically destabilised by the loss of cell-to-cell contact on the injured side, and this stress triggers a specific transcriptional response of the border zone myocardium. *Ankrd1*, being a mechanotransduction and stress-responsive gene upregulated after cardiac injury (42, 63), may participate in this transcriptional response, and presumably be able to sense and transduce these mechanical stress signals.

### Limitations of the study

Transcriptomics analysis was performed only at 7 dpci. Still, our results are suggesting that transcriptome profiling of the *ankrd1a* mutant at earlier and later time points could provide more information in regard to the mechanism and regulation of its early activation, potential implication in immune response, and the role in scar removal and functional recovery. Another limitation is the lack of a physiological assessment of cardiac function after mutant heart healing.

## CONCLUSIONS

The activity of *ankrd1a* is associated with the processes occurring in the border zone CMs and dependent on the level of scar resolution, supporting the presumed role of *ankrd1a* in protecting CMs from mechanical damage on the border with non-contractile tissue. Results presented in this manuscript set the stage for the next step in deciphering the presumed function of *ankrd1a* in immune response in the injured zebrafish heart. Taking into account new information on *ankrd1a* expression and function in skeletal muscle repair (45) and cardiac regeneration (this study) in zebrafish, we may consider *ankrd1a* more as a marker and fine-tuner, rather than an essential player during the healing of injured striated muscle.

## SUPPLEMENTAL INFORMATION

Figures S1 and S2, Tables S1 and S2 (https://doi.org/10.6084/m9.figshare.29376512) Supplemental Table S3 (https://doi.org/10.6084/m9.figshare.29376506)

Supplemental Table S4 (https://doi.org/10.6084/m9.figshare.29377241) Supplemental Table S5 (https://doi.org/10.6084/m9.figshare.29377262)

## ACKNOWLEDGMENTS

We are grateful to dr Filipa Simões for the insightful discussion of the results and helpful comments.

## DATA AVAILABILITY

The RNA-seq data reported in this manuscript have been deposited in the Gene Expression Omnibus (GEO) database with the accession number GSE285387. Other relevant data can be found within the article and supplementary information.

